# Overlooked biodiversity loss in Amazonian smallholder agriculture

**DOI:** 10.1101/192955

**Authors:** Jacob B. Socolar, Elvis H. Valderrama Sandoval, David S. Wilcove

## Abstract

Diversified smallholder agriculture is the main human land-use affecting the western Amazon, home to the world’s richest terrestrial biota, but the scant available data to date have suggested that the biodiversity impacts of this land-use are small. Here, we present comprehensive surveys of birds and trees in primary forest and smallholder agricultural mosaics in northern Peru. These surveys reveal substantial biodiversity losses that have been overlooked by other studies. Avian biodiversity losses arise primarily from biotic homogenization across infrequently surveyed forest habitats (a loss of beta-diversity). Furthermore, tree species richness declines much more steeply than bird richness. Statistical modeling of local habitat features that allow forest-associated species to persist in the smallholder mosaic strongly suggests that our results represent a best-case scenario for Amazonian agricultural biodiversity. We conclude that previous assessments of the biodiversity value of Amazonian smallholder agriculture have been overly optimistic because they are restricted to upland habitat, thereby missing losses in beta diversity; do not evaluate trees; and/or rely on generalizations from less speciose areas of the Neotropics, where habitat specialization amongst species is less prevalent. Smallholder agriculture will likely expand in western Amazonia due to infrastructure development, and it must be seen as a serious threat to the region’s biodiversity.

## INTRODUCTION

The western Amazon is the global epicenter of terrestrial biodiversity (Jenkins et al. 2013) and the largest remaining tropical forest wilderness (Tyukavina et al. 2015), but it is nevertheless threatened by human activities. In contrast to the mechanized agriculture and ranching in southeastern Amazonia, the principal driver of forest loss in the western Amazon is smallholder slash-and-burn agriculture (Finer & Novoa 2016; Ravikumar et al. 2017). This practice creates mosaics of cultivations and secondary forest surrounding human settlements. The prospect of increased smallholder settlement in western Amazonia in the wake of roadbuilding and hydrocarbons development has raised alarm for this bastion of tropical biodiversity. For example, most of western Amazonia is covered in hydrocarbons concessions, the development of which would provide road access for settlers (Finer & Orta-Martínez 2010; Laurance et al. 2014).

Existing data on biodiversity in western Amazonian agriculture are extremely scant, consisting of two small-scale studies of birds and dung beetles, respectively (Andrade & Torgler 1994; Korasaki et al. 2013). These studies document comparable avian richness in slash-and-bum mosaic and primary forest (Andrade & Torgler 1994), and comparable dung beetle richness in young secondary forest and primary forest (Korasaki et al. 2013). These isolated results contrast with results obtained from more intensive land-use change in the eastern Amazon (e.g. large-scale agriculture, fragmentation, silviculture, or fire; Ferraz et al. 2007; Barlow et al. 2007; Berry et al.2008; Mahood et al. 2011; Gardner et al. 2013; Lees et al. 2015; Moura et al. 2016). However, they agree with numerous Mesoamerican studies that have documented high levels of biodiversity in smallholder mosaics (Daily et al. 2001; Sekercioglu et al. 2007; Ranganathan et al. 2008; Karp et al. 2011; Mendenhall et al. 2011) and have generated sustained debates over the relative conservation benefits of protecting primary-forest versus preventing agricultural intensification/industrialization of smallholder mosaics, especially given limited funding for conservation (Phalan et al. 2011; Gibson et al. 2011). This debate has a special urgency in the western Amazon, where forests remain largely intact but under increasing threat from smallholder agriculture, including inside protected areas (Finer & Novoa 2016).

Despite concordant results from Andrade and Torgler (1994) and Korasaki et al. (2013) that would downplay the significance of smallholder agriculture and conversion of primary forest to secondary forest, there are strong reasons to think that the impacts of smallholder activities on Amazonian biodiversity might be more severe than previously recognized. Thus, the biodiversity impacts of the main land-use affecting the world’s richest terrestrial biota remain unknown. First, the few Amazonian studies that have examined smallholder agriculture either have included relatively few smallholder sites (e.g. 7 out of 361 sites in smallholder habitats in Moura et al. 2013) or are limited in their total sampling (Andrade & Torgler 1994; Korasaki et al. 2013). For example, Andrade and Torgler ( 1994) found bird diversity comparable to primary forest in Colombian slash-and-burn mosaics, but this conclusion rests on only understory birds sampled over a relatively small area.

Second, Amazonia is more species-rich than other areas of the Neotropics, so data from Mesoamerica might not generalize to the Amazon. Ecological theory predicts that habitat specialization among species should be more frequent in hyperdiverse communities (MacArthur & Levins 1967), and this might predispose Amazonian communities to be more sensitive to habitat alteration. Consistent with this idea, modelling work suggests that a given deforestation scenario would impact Amazonian bird communities more heavily than their Mesoameiican counterparts (Newbold et al. 2014).

Third, previous studies focused on the upland (*terra firme*) forest of uplifted clay terraces. Yet Amazonia contains additional forest types that are critical for biodiversity and are also impacted by slash-and-burn. These include floodplain habitats, bamboo forests, and forests on white-sand soils, all of which harbor specialist species that do not occur in terra firme forests (Remsen & Parker 1983; Wittmann et al. 2006; Fine et al. 2010; Alvarez Alonso et al. 2013; Socolar et al. 2013). Because biotic homogenization can drive landscape-scale biodiversity loss in tropical forests (Karp et al. 2012; Solar et al. 2015; Alroy 2017; Giam 2017), effective conservation planning requires an extensive comparison of biodiversity in intact and degraded landscapes across multiple forest types (Socolar et al. 2016). However, we are unaware of any data from the western Amazon that evaluate the biodiversity consequences of land-use change across multiple forest-types simultaneously.

Here, we quantify the biodiversity consequences of Amazonian slash-and-bum agriculture based on extensive field surveys of bird and tree diversity in Loreto department, Peru. In Loreto, upland, floodplain, and white-sand forests collectively harbor the richest avifauna and tree flora on Earth (ter Steege et al. 2013). Although the area remains largely roadless, the city of Iquitos is the world’s largest city without an outside road link (circa 0.5 million inhabitants), and slash-and-bum mosaics are ubiquitous along rivers and local roads (Mäki et al. 2001). Furthermore, slash-and-bum is practiced to varying degrees, often legally, inside the region’s protected areas (Pulgar Vidal & Gamboa Moquillaza 2013). Therefore, our results stand to inform conservation practices and priorities across this hyperdiverse yet understudied region in the face of ongoing smallholder expansion.

Despite previous work suggesting that the biodiversity value of tropical smallholder landscapes is very high (Sekercioglu et al. 2007; Karp et al. 2011; Mendenhall et al. 2011), we hypothesized that habitat specialists would fare poorly at disturbed sites, driving landscape-scale biodiversity declines via a reduction in beta-diversity (Karp et al. 2012; Socolar et al. 2016).

## METHODS

### STUDY SITES

We conducted fieldwork in the Amazonian lowlands of Loreto Department, Peru within 230 km of the city of Iquitos. Natural habitats in the region are varied and interdigitate at fine spatial scales. We focused on four terrestrial habitats that harbor distinctive biological communities. Quintessential *upland forest* grows on uplifted clay-soil terraces of (Higgins et al. 2011). These uplands are the most spatially extensive habitat in the region and the richest in bird and tree species.

*Floodplain forest* along major rivers, subject to protracted flooding during January-June (Espinoza et al. 2013), differs from the uplands in tree and bird species composition (Remsen & Parker 1983; Wittmann et al. 2004). *White-sand forest* occurs on deposits of pure white-sand soil (arenosols) and supports a characteristic avifauna and flora that is absent from other habitats (Fine et al. 2010; Alvarez Alonso et al. 2013). Lastly, *river islands* harbor *Cecropia* (Urticaceae)-dominated woodland with a characteristic suite of specialist birds (Rosenberg 1990). Slash-and-burn agriculture affects all of these habitats, removing primary forest vegetation and replacing it with a heterogeneous mosaic of clearings, hedgerows, and secondary forests (Figure 1). Typical crops include manioc, corn, camu-camu, and watermelon on floodplains; manioc, plantain, rice, small buffalo pastures, and small aquaculture ponds in uplands; manioc and pineapple on white sands; and rice, watermelon, and manioc on islands.

**Figure 1:**
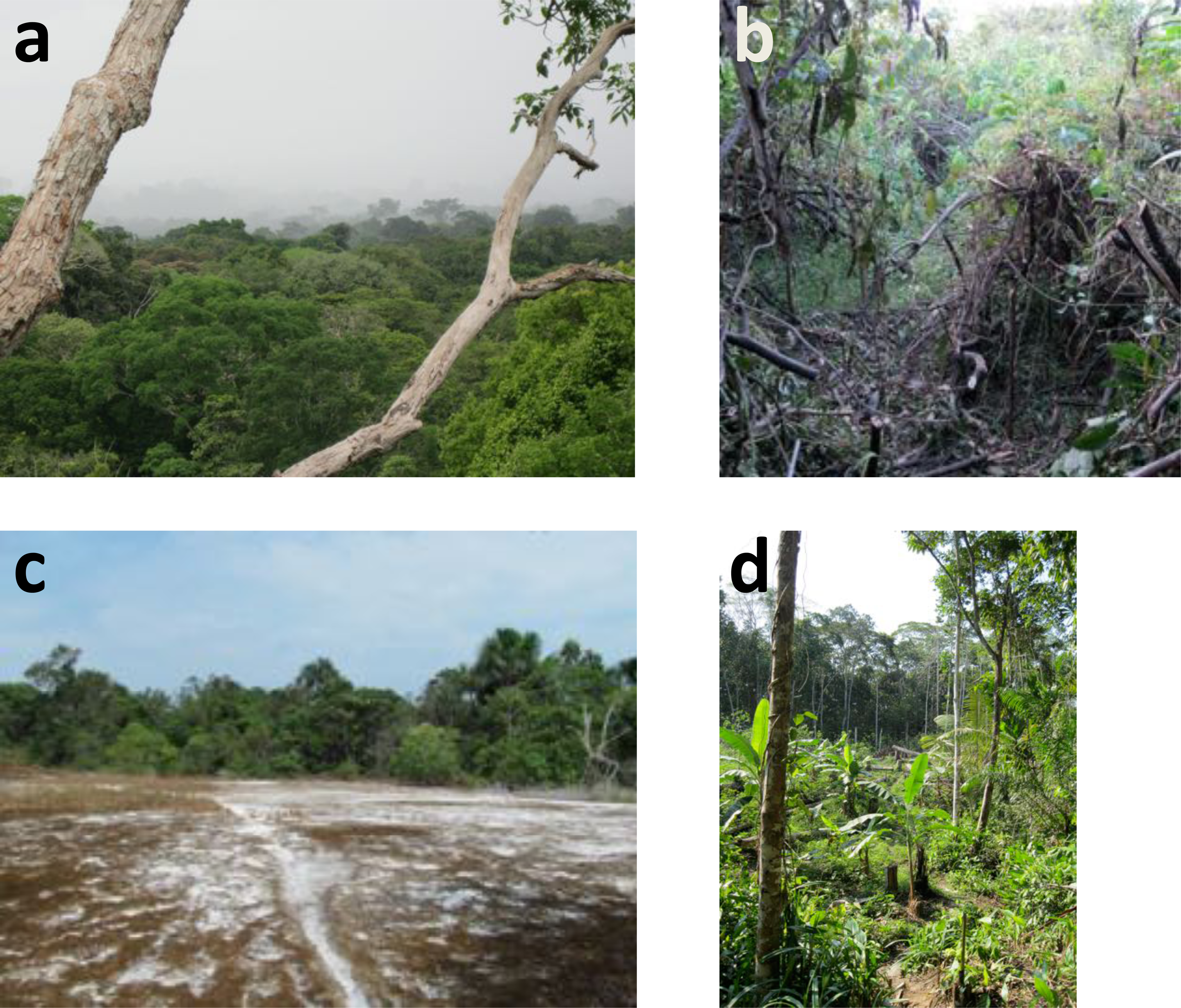
In western Amazonia, slash-and-burn agriculture converts primary forest **(a)** to various disturbed habitats across uplands, floodplains, and white-sands, resulting in a heterogeneous mosaic of secondary habitats. Shown here are tangles following abandonment of a floodplain agricultural plot **(b)**, barren ground and scrub following agricultural abandonment on white-sands **(c)**, and a mosaic of upland secondary forest and active agricultural plots **(d)**. **(b-d)** represent the range of slash-and-burn habitats in a highly diversified mosaic, not typical differences between soil types.

We sampled bird and tree communities at intact sites (primary forest) and disturbed sites (slash-and-burn mosaics of active cultivation and fallow secondary forest). We selected twenty intact sites within 230 km of Iquitos harboring accessible habitat that is largely undisturbed by humans for as long as records are available, except for light selective logging at floodplain sites and widespread hunting of game animals (see supplementary material). These sites spanned the major forest habitats of the region: ten in uplands spanning both banks of the Amazon River, six on floodplains, and four in white-sand forest. We were unable to find intact examples of river islands large enough to accommodate our sampling scheme. We then selected twenty disturbed sites in slash-and-bum mosaic, each paired with an intact site for forest type, soil texture, and geographic proximity. At each study site, we established six sampling points spaced by at least 210 meters to avoid double-counting during avian point counts.

During subsequent vegetation assessment, we determined that six sampling points on different transects were unsuitable for analysis due to their inadvertent location in transitional habitat at the edge of the forest-type of interest. Wooding and time constraints prevented us from sampling trees at two study sites (one in intact floodplain and another in intact uplands), and we removed their paired disturbed sites from the tree dataset. Thus, the final dataset contained 234 bird sampling points and 209 tree sampling points. See supplementary material for details of site selection, site spacing, and site characteristics.

### BIODIVERSITY DATA

We surveyed birds and trees at each sampling point. For birds, a single observer (REDACTED) conducted four ten-minute 100-meter-radius point-counts at each sampling point during july-December 2013-2014. Surveys ran from first light until mid-morning, and were not conducted in rain or windy conditions. We visited most points in both years and rotated the visit order to ensure that each point received early-morning coverage. To assemble our final dataset for analysis, we aggregated data across the four visits to each point by taking the maximum count for each species from any visit.

We made two modifications to standard point-count protocols, tailored to the challenges of detecting skittish species and birds in mixed-species flocks (see supplementary material). First, we included detections of species that flushed during our approach to and departure from each point (within 100 m). Second, when mixed flocks detected during the point count lingered within 100 meters after the count period, we proceeded to follow the flock until we identified all of its participants or until it moved >100 m from the point. We separately recorded individuals detected via these modifications, permitting us to include them or exclude them from analysis (see SENSITIVITY ANALYSIS, below).

To survey trees, we established a 50x2 m^2^ tree plot at a fully randomized location within 100 m of each sampling point (equivalent to 0.6 Gentry transects per site; (Gentry 1988). Within these plots, we identified ever) tree greater than 2.5 cm diameter at breast height. We collected a voucher for each species (except for palms with very large leaves), deposited in the herbarium at the Universidad Nacional de la Amazonía Peruana (UNAP). One botanist (REDACTED) conducted all sampling and made all species determinations with reference to the UNAP herbarium collections. See supplementary information for detailed bird and tree survey protocols.

### BIODIVERSITY COMPARISONS

We used sample-based rarefaction to compare bird and tree richness in intact and disturbed landscapes on a per-area basis (Chao et al. 2014). For trees, this revealed dramatic diversity loss due to massively reduced densities of individuals at disturbed sites (i.e., cleared areas have fewer trees). Therefore, we used individual-based rarefaction to test for a second-order effect of slash-and-burn on tree diversity, controlling for the number of individuals sampled. For both birds and trees, we performed rarefaction analysis on each forest type separately (upland, floodplain, white-sand) and for all forest types combined. We also visualized patterns of community change using non-metric multidimensional scaling.

Some bird species that we did not record at intact sites are well known to be common on intact river-islands (Rosenberg 1990). As noted above, we were unable to sample intact river island habitat because in our study area all accessible river islands large enough to accommodate our sampling scheme have been settled, cleared, or otherwise disturbed by people. Therefore, we conducted a follow-up analysis to account for bias related to the intact river-island avifauna. We obtained a comprehensive list of bird species that were common on intact river-islands within the study area thirty years ago (Rosenberg 1990). We then repeated our analysis while excluding these species from all datasets, thereby removing their influence on our conclusions. We stress that we selected these species not because they are prevalent in disturbed samples, but because they are known to be prevalent in an intact river-island habitat that we were unable to sample. By removing only common river-island species, we are confident that we removed very few species that would not have appeared in the dataset for intact forest types, had we been able to sample river islands. Therefore, this analysis mitigates bias in the comparison between intact and disturbed habitats.

### POPULATION COMPARISONS

For every species of bird and tree in the dataset, we calculated Bayesian point-estimates and 95% credible intervals for the multiplicative change (fold-change) in abundance between intact and disturbed sites. To do so, we assumed that the number of individuals detected at intact and disturbed sites were realizations of Poisson processes. This implies that the total count at disturbed sites is a binomial draw from the summed count at intact and disturbed sites, and furthermore that the logarithm of the fold-change between the Poisson means is equal to the logit of the binomial proportion *p* (Przyborowski & Wilenski 1940). We computed the posterior density of *p* using the Jeffreys prior, and we used the posterior density of *p*/(1-*p*) for inference on the fold-change (Brown et al. 2001).

### DISTRIBUTION OF DISTURBANCE-SENSITIVE SPECIES

To understand what features of disturbed points allow them to support species characteristic of intact forests, we defined *disturbance-sensitive species* as those that are more abundant in intact forest than disturbed forest, and *disturbance-sensitive counts* as the total number of individuals belonging to disturbance-sensitive species detected at each point. We then fit generalized linear mixed models for birds and trees to assess the relationship between disturbance-sensitive counts and local habitat data (see below) across the disturbed points.

#### Local habitat data

At every sampling point, we recorded the number and size of streams and estimates of percent cover of 10 vegetation formations within 100 m of the point (see supplementary material). Using Landsat 8 imagery downloaded from the Global Forest Change Data website (Hansen et al 2013), we built a random-forest classifier of the study landscape as intact, disturbed, or open water at 30 m resolution. We validated our classification against the central coordinates of our 240 sampling points, and then we extracted the area classified as intact within 200, 500, and 5000 m of each disturbed sampling point. We also measured the distance from each disturbed point to the nearest primary forest (continuously forested since 1985, before the acceleration of forest clearance in the region; (Mäki et al. 2001) and to the nearest river (channel width > 30 m) based on visual examination of Landsat imagery in the USGS Landsat Look viewer, supplemented with aerial imager in Google Larth.

#### Mixed models

Initially, we assumed that any species recorded in higher numbers in intact than disturbed habitat is disturbance-sensitive. I^1^’or birds and trees, we fit ordinary and zero-inflated Poisson and negative binomial mixed models (treating study-site as a random effect) for the disturbance-sensitive counts using a variety of predictors describing local vegetation cover at the 100 m scale, forest cover at 0.2 – 5 km spatial scales, and proximity to major rivers (see supplementary material). We used the small-sample corrected Akaike information criterion (AICc) to select covariates and error structure that yielded parsimonious models, and we base inference on broad agreement across all top-performing models.

To verify that our conclusions were robust to uncertainty in which species are disturbance-sensitive, we re-analyzed the model with the lowest AICc score as follows. Using the binomial likelihood described above, we computed the probability that each species in the dataset is disturbance-sensitive by integrating the posterior distribution for the binomial proportion (based on a uniform prior) from 0 to 0.5. We then randomly assigned each species to be disturbance-sensitive or not based on these probabilities, re-computed the disturbance-sensitive counts, and fit the regression model to these counts under a Bayesian mode of inference using Markov-chain Monte Carlo sampling implemented in JAGS (Plummer 2003). We repeated this process 500 times, combined the posterior chains for inference, and compared the resulting parameter estimates to the corresponding frequentist estimates.

### SENSITIVITY AND DETECTABILITY’ ANALYSIS FOR BIRDS

To ensure that our non-standard point-count methodology did not bias avian sampling, we repeated our analyses using only detections obtained via standard protocols. We used an N-mixture model to determined that avian detectability is likely to be at least as high in disturbed habitats as intact habitats (see supplementary material). Therefore, if anything, our results overestimate the biodiversity value of smallholder landscapes.

## RESULTS

Across pristine and disturbed habitats combined we recorded 455 bird species and 751 tree species; the bird dataset is among the richest single-observer point-count datasets ever assembled. We found very high avian richness in slash-and-burn mosaics. In fact, in each habitat studied (uplands, floodplain, white-sands), sample-based rarefaction revealed that bird richness at disturbed sites was comparable to intact sites (Figure 2). However, tree richness declined severely. This decline partly resulted from dramatic reductions in the number of individuals at disturbed sites (i.e. cleared areas have fewer trees) but was exacerbated by changes in the species-abundance distribution (Figure 2).

**Figure 2:**
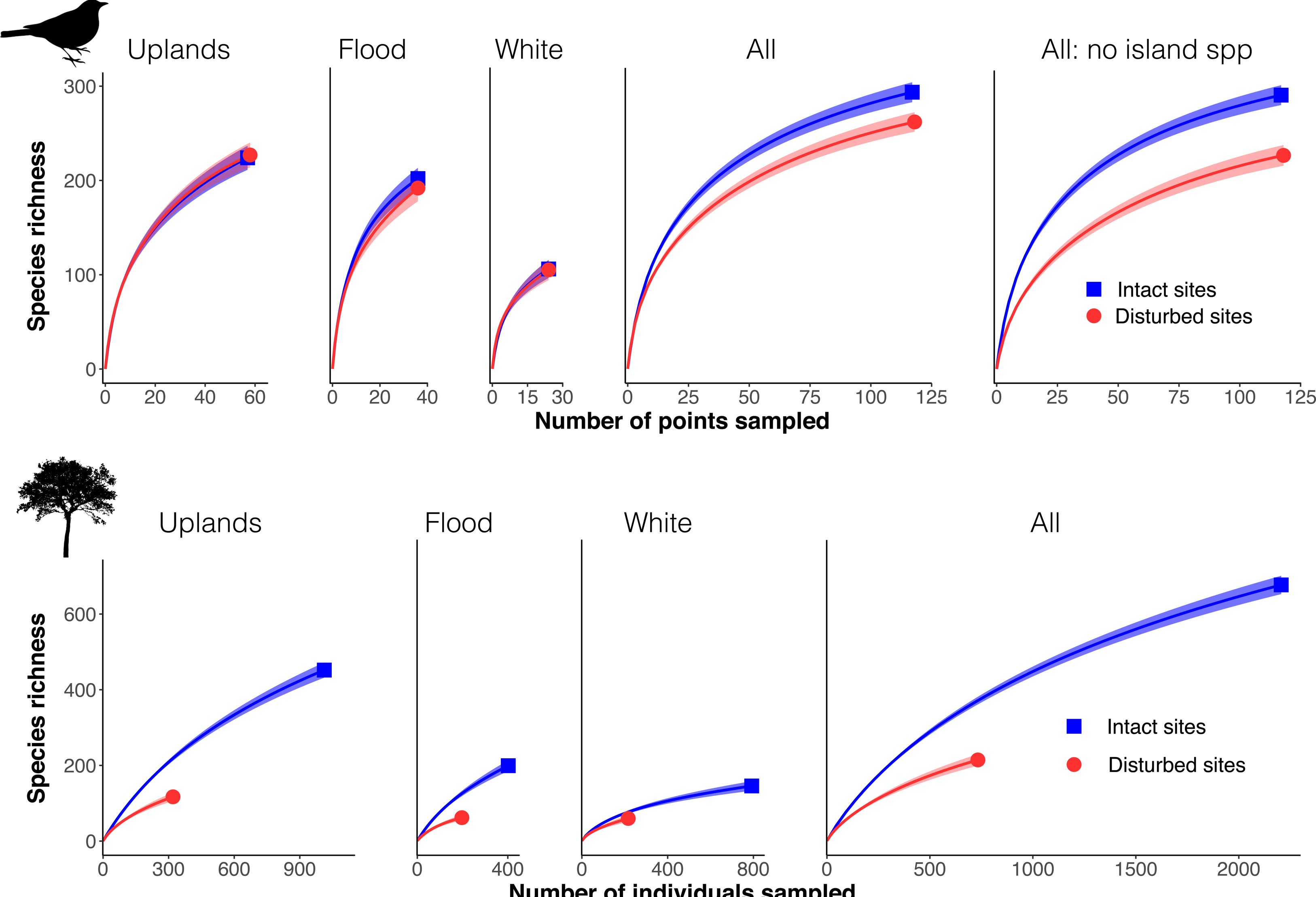
Sample-based rarefaction (mean and 95% confidence interval) for birds (top row) shows that within each forest type, disturbed sites are as species-rich as their intact counterparts. However, when forest-types are aggregated, intact forest is more diverse, especially after accounting for the distribution of river-island species. Lor trees (bottom row), individual-based rarefaction shows that richness plummets in disturbed forests, due to low individual abundance and a second-order effect of altered species-abundance distributions after controlling for the number of individuals sampled. Aggregated forest-types do not show greater tree-richness differences than individual forest-types; instead, the uplands dominate the species pool and are effectively as diverse as all habitats combined.

Importantly, considering each habitat in isolation substantially underestimated the difference in bird richness between intact and disturbed landscapes. Across habitats, reductions in beta-diversity caused modest but significant declines in bird richness. Moreover, the apparent biodiversity value of smallholder landscapes was substantially inflated by the spurious absence of river-island species from our intact sites (an artifact of our inability to sample intact river islands). When the influence of these poorly sampled river-island species is removed from both the intact and disturbed points, it becomes apparent that intact landscapes have dramatically higher avian richness than disturbed landscapes in our study region (Figure 2). This occurs because the river-island avifauna overlaps more with disturbed habitats than with other intact habitats in the study area. We did not observe a similar pattern in trees, though non-metric multidimensional scaling suggests that some homogenization might have occurred (Figure 3). Instead, uplands dominated the tree species richness of all intact sites combined, minimizing the opportunity for specialists in other habitats to contribute to richness patterns (Figure 2).

**Figure 3:**
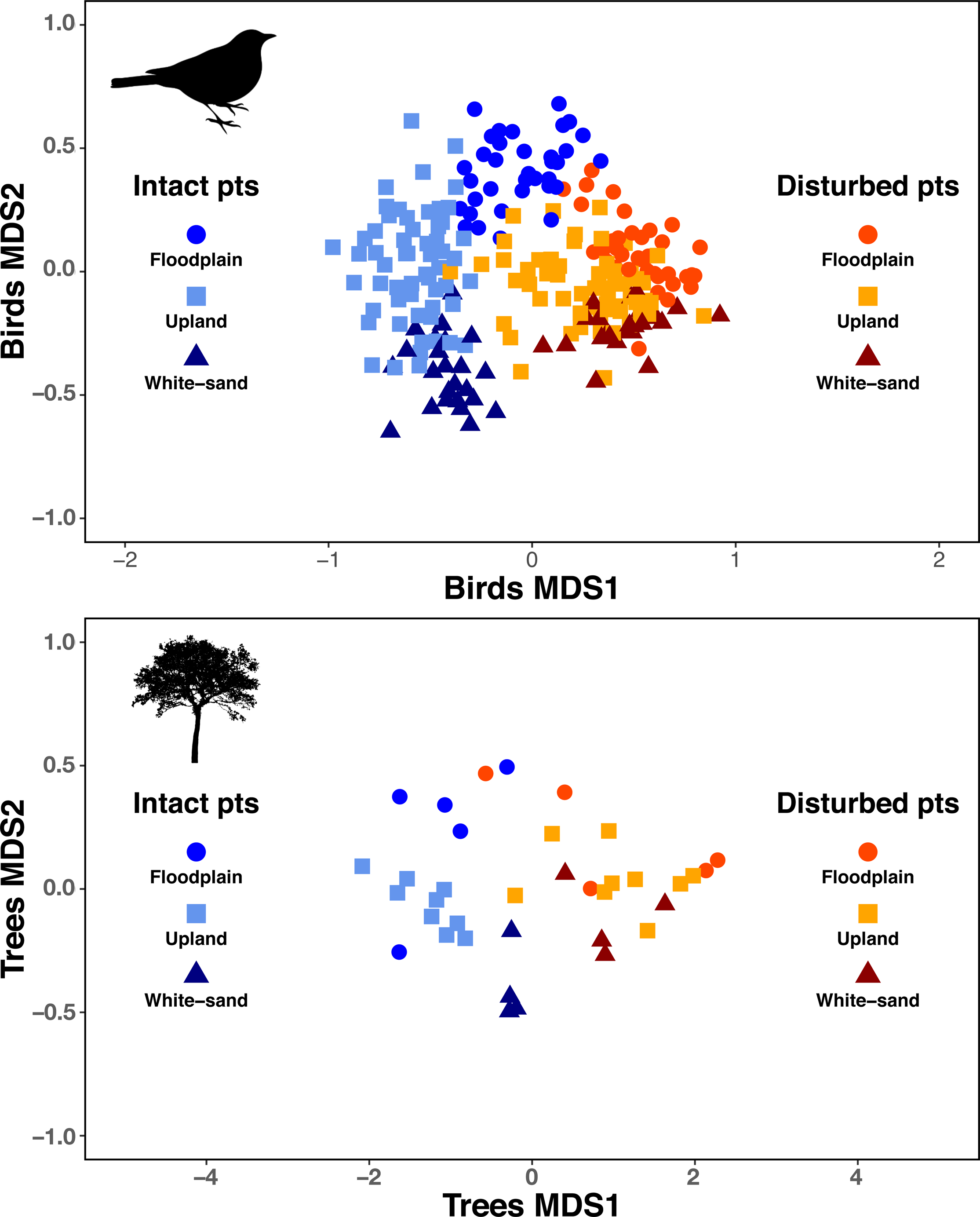
Non-metric multidimensional scaling (NMDS) based on Raup-Crick dissimilarities for point-scale bird data (top; stress = 0.21) and site-scale tree data (bottom; stress = 0.22; point-scale tree data were too sparse for NMDS). In both cases, the first NMDS axis captures the difference between intact and disturbed sites, while the second axis captures the gradient from nutrient-rich floodplains to nutrient-poor white-sands. Intact and disturbed sites segregate almost completely. Heterogeneity between forest types at intact sites is collapsed at disturbed sites.

Disturbed sites consistently clustered separately from intact sites in terms of their species composition, and non-metric multidimensional scaling of community composition revealed that the difference between intact and disturbed sites corresponded to the first axis of variation (Figure 3). The second axis of variation, corresponding to an edaphic gradient from floodplains through uplands to white sands, was collapsed at disturbed sites, reflecting the loss of beta-diversity among forest types. These patterns are consistent for birds and trees and for a variety of incidence- and abundance-based dissimilarity metrics (Figure S5). Thus, disturbance in addition to driving species loss, smallholder agriculture drives the disassembly and re-arrangement of primary forest bird communities.

Furthermore, large numbers of disturbance-sensitive species showed dramatically reduced abundance at disturbed sites (Figure 4). For example, we detected the Screaming Piha (*Lipaugus vociferans*) 137 times at intact sites, and only once at disturbed sites. Similarly, we detected the tree *Eschweilera coriacea* (Lecythidaceae) thirty-one times at intact sites and only once at disturbed sites. In the rarefaction analysis, such species contribute to the disturbed-site total, but in fact they are severely harmed by slash-and-burn practices. Among the 249 bird and 221 tree species for which we detected a significant change in abundance, 57% and 86% declined, respectively. Of the birds that significantly increased in abundance in our dataset, fully 39% are common on intact river islands (Rosenberg 1990). Failure to detect significant abundance changes was generally a consequence of low sample size (and probably not a consequence of small effect size). The median sample size among species without a significant effect was two for birds and one for trees.

**Figure 4:**
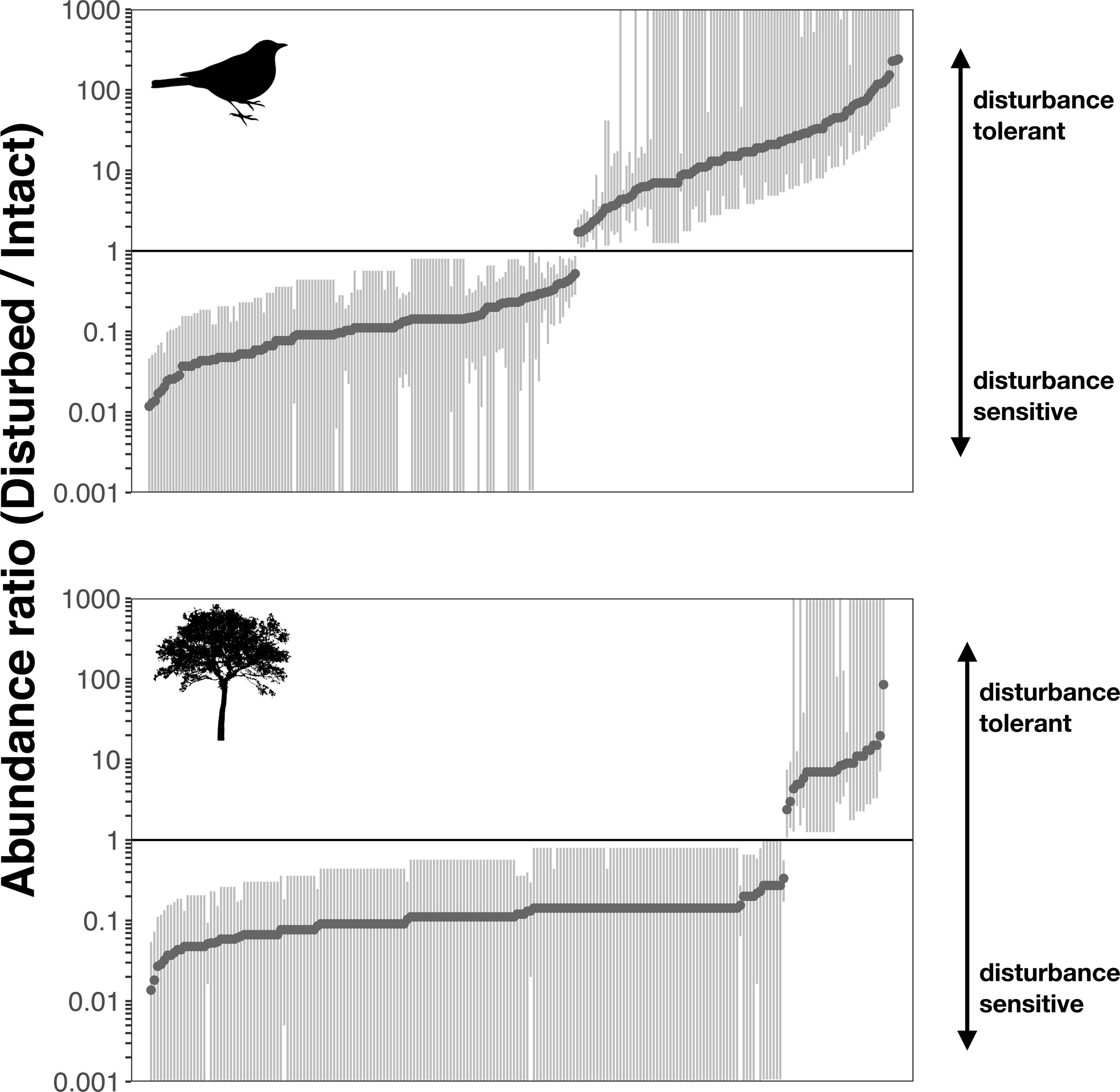
Multiplicative changes in abundance for birds (top) and trees (bottom). Species significantly different from one (i.e. abundance differs by land-use class) are given by dark points with 95% credible intervals. Most tree species plummet in abundance. Bird communities include species that fare well following disturbance, but 57% of species with significant abundance-changes declined, often dramatically (note the logarithmic y-axis).

Mixed models revealed a major positive influence of local forest cover and nearby primary forest on the abundance of disturbance-sensitive birds and trees that was consistent across all well-performing models (Table 1). For birds, the most important components of this effect were primary forest cover at a radius of 5 km and secondary forest cover at a radius of 100 m. For trees, the key components were secondary forest cover at a radius of 100 m and primary forest cover at a radius of 200 m. These effects were robust despite uncertainty in which species are disturbance-sensitive.

**Table 1:**
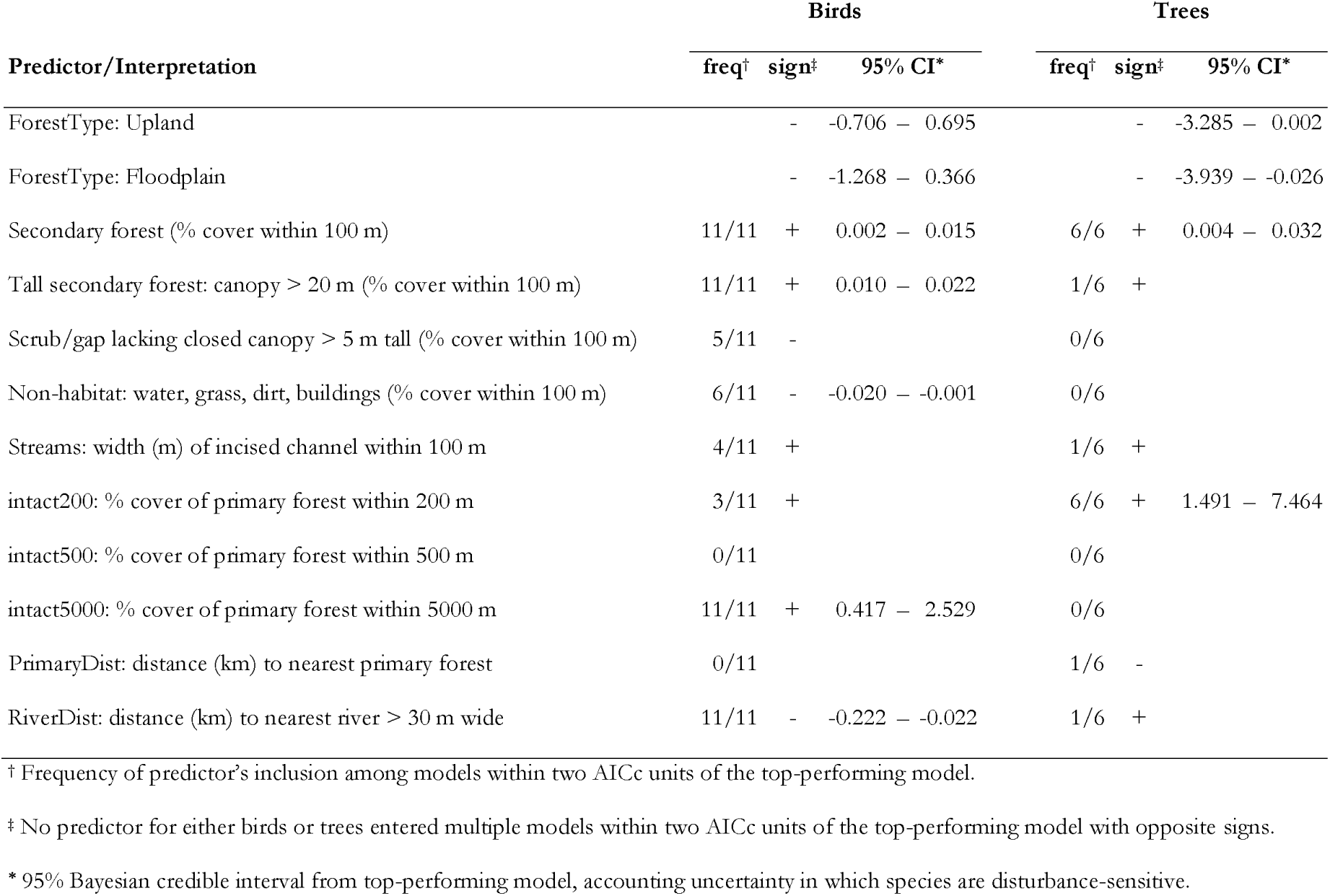
Results of models for counts of disturbance-sensitive birds and trees, summarizing results for the top performing model (credible intervals for the effect size) and for all models within 2 AICc units of the top performing model (frequency of inclusion and sign of effect). Forest type was included in all models as a control. All of the best-performing bird models used a NB1 negative binomial error structure without zero-inflation. Five of the best-performing tree models (including the top model) used a zero-inflated NB2 negative binomial error structure, one used NB2 error without zero-inflation.

## DISCUSSION

Our results constitute the first large-scale biodiversity assessment of slash-and-bum agriculture in western Amazonia, and one of the first biodiversity assessments in degraded Amazonian landscapes to explicitly consider multiple natural habitat types. These features define a key knowledge gap for conservation science, because western Amazonia is the epicenter of terrestrial biodiversity on Earth (Jenkins et al. 2013), harbors multiple types of forest, is heavily affected by slash-and-burn agriculture (Finer & Novoa 2016), and features extensive species turnover (beta-diversity) between natural habitats (Tuomisto et al. 1995; Pomara et al. 2012).

Our results are sobering. Diversity loss, community turnover, and large numbers of declining, disturbance-sensitive species characterize the transition from intact forest to slash-and-burn mosaic. Slash-and-burn agriculture collapses avian beta-diversity across forest-types, and this process drives substantial reductions in gamma-diversity for birds. Because slash-and-bum mosaics are as diverse as primary forest within any single forest type (e.g. within upland foreest), previous work was unable to detect this decline (Andrade & Torgler 1994; Korasaki et al. 2013). Previous studies have treated an environmental domain equivalent to the first panel of our Figure 2.

Moreover, within the slash-and-burn mosaic, secondary forest cover and proximity to primary forest were consistent, strong predictors of the occurrence of disturbance-sensitive species. According to our vegetation classifier, the median disturbed point in our dataset was surrounded by over 19% primary forest at a radius of 0.2 km, 28% at 0.5 km, and 57 % at 5 km. The proximity of intact habitat, coupled with the high heterogeneity and low land-use intensity of the slash-and-bum mosaic (the median disturbed point contained 30% closed-canopy secondary-forest cover within a 100 m radius), strongly suggests that our results are a best-case scenario for biodiversity in Amazonian smallholder agriculture. The conservation value of slash-and-burn mosaics in our study area depends on extensive fallow areas (i.e. secondary forests) and spillover from primary forest.

Recent work from elsewhere in Amazonia suggests that the biodiversity impacts of smallholder agriculture might be even more severe than our methods can detect. Space-for-time substitutions might underestimate the severity of impacts in Amazonia (Farança et al. 2016), perhaps due to inadequate primary-forest baseline data. Furthermore, the negative impacts of agricultural disturbance can spill across into adjacent primary forest, leading to substantial additional losses of conservation value (Barlow et al. 2016). In our study area, a few species (e.g. curassows in the genus *Mitu*) are so heavily hunted that they are absent even at intact sites, and our analyses cannot shed light on their disturbance-sensitivity.

Some implications of our results extend beyond the western Amazon. In particular, we note that many previous comparisons of biodiversity value at intact and degraded tropical sites have been restricted to a single natural habitat (or have analyzed multiple habitats separately), with variable results (e.g Daily et al. 2003; Peh et al. 2006; Ranganathan et al. 2008; Phalan et al. 2011; Kurz et al. 2014). Our results show that the large-scale pattern across multiple habitats is gloomier than single-habitat results suggest, at least for birds. This conclusion is consistent with the observation that smallholder agriculture reduces pairwise avian compositional dissimilarities across biogeographic regions of Costa Rica (Karp *et al.* 2012). We expand on this result by showing that the homogenization produced by smallholder agriculture drives substantial losses of regional gamma-diversity (this is not a forgone conclusion; see Socolar *et al.* 2016). Moreover, we show that homogenizing effects are important not only across widely spaced biogeographic regions, but also across fine-scale habitat formations that structure Amazonian communities. The vast majority of biodiversity assessments of Neotropical agriculture have focused on uplands and therefore missed the additional biodiversity losses driven by homogenization across forest-types. Habitat differences within biogeographic regions are globally ubiquitous (e.g. due to variation in elevation, soils, hydrology, climate, etc), and revealing the full impacts of disturbance requires sampling that is both spatially extensive and locally comprehensive with respect to habitat variation (Gardner et al. 2013; Solar et al. 2015). Habitat specialization and spatial turnover are characteristic of hyperdiverse species communities, suggesting that habitat degradation might have its worst effects precisely where biodiversity is highest.

We also note that previous studies of biodiversity in Neotropical agricultural landscapes have broadly neglected trees. A recent meta-analysis of the biodiversity value of degraded tropical landscapes was unable to include a single study of tree diversity in Neotropical agriculture (Gibson et al. 2011). This situation might arise because lower tree diversity in cleared areas is perceived as a forgone conclusion (more attention has been paid to shrubs and forbs, e.g. (Mayfield & Daily 2005). Nevertheless, trees make up a critical component of tropical biodiversity, and maintaining tropical tree diversity is likely essential for the long-term conservation a variety of coevolved species (Koh et al. 2004). Moreover, the impacts of agriculture on tree diversity are even more severe than could be predicted by declines in abundance alone; agricultural landscapes are species-poor even after controlling for the number of individual trees sampled. Thus, field inventories of tree communities are crucial for accurately assessing the biodiversity consequences of slash-and-burn agriculture, and our results paint a bleak picture.

We do not mean to dismiss innovative efforts, including efforts inside protected areas, to harmonize conservation objectives with the livelihoods of local people (Pulgar Vidal & Gamboa Moquillaza 2013). There is a clear humanitarian mandate for such efforts, and they can prevent the even greater losses of biodiversity that result from the conversion of disturbed forests and agricultural mosaics to soy monocultures or tree plantations. However, we do mean to sound the alarm over the potential consequences of ongoing smallholder expansion. There will be severe biodiversity losses if settlers gain access to the last remaining tropical wildernesses in western Amazonia, no matter how lightly they tread.

## ACKNOWLEDGMENTS

REDACTED FOR REVIEW.

## REFERENCES

Alroy J. 2017. Effects of habitat disturbance on tropical forest biodiversity. Proceedings of the National Academy of Sciences 114:6056–6061.

Alvarez Alonso J, Metz MR, Fine PV. 2013. Habitat specialization by birds in western Amazonian white-sand forests. Biotropica 45:365–372. Wiley Online Library.

Andrade GI, Torgler HR. 1994. Sustainable use of the tropical rain forest: evidence from the avifauna in a shifting-cultivation habitat mosaic in the Colombian Amazon. Conservation Biology 8:545–554.

Barlow J et al. 2007. Quantifying the biodiversity value of tropical primary, secondary, and plantation forests. Proceedings of the National Academy of Sciences of the United States of America 104:18555–18560.

Barlow J et al. 2016. Anthropogenic disturbance in tropical forests can double biodiversity loss from deforestation. Nature 535:144–147. Nature Publishing Group.

Berry NJ, Phillips OL, Ong RC, Hamer KC. 2008. Impacts of selective logging on tree diversity across a rainforest landscape: the importance of spatial scale. Landscape Ecology 23:915–929.

Brown LD, Cai TT, DasGupta A. 2001. Interval estimation for a binomial proportion. Statistical science 16:101–117.

Chao A, Gotelli NJ, Hsieh TC, Sander EL, Ma KH, Colwell RK, Ellison AM. 2014. Rarefaction and extrapolation with Hill numbers: a framework for sampling and estimation in species diversity studies. Ecological Monographs 84:45–67.

Daily GC, Ceballos G, Pacheco J, Suzán G, Sánchez-Azofeifa A. 2003. Countryside biogeography of neotropical mammals: conservation opportunities in agricultural landscapes of Costa Rica. Conservation Biology 17:1814–1826. Wiley Online Library.

Daily GC, Ehrlich PR, Sanchez-Azofeifa A. 2001. Countryside biogeography: use of human-dominated habitats by the avifauna of southern Costa Rica. Ecological Applications 11:1–13.

Espinoza JC, Ronchail J, Frappart F, Lavado W, Santini W, Guyot JL. 2013. The major floods in the Amazonas river and tributaries (western Amazon basin) during the 1970—2012 period: a focus on the 2012 flood. Journal of Hydrometeorology 14:1000–1008.

Ferraz G, Nichols JD, Hines JE, Stouffer PC, Bierregaard RO, Lovejoy TE. 2007. A large-scale deforestation experiment: effects of patch area and isolation on Amazon birds. Science 315:238–241.

Fine PVA, García-Villacorta R, Pitman NCA, Mesones I, Kembel SW. 2010. A floristic study of the white-sand forests of Peru. Annals of the Missouri Botanical Garden 97:283–305.

Finer M, Novoa S. 2016. Large-scale vs. small-scale deforestation in the Peruvian Amazon. MAAP 32.

Finer M, Orta-Martínez M. 2010. A second hydrocarbon boom threatens the Peruvian Amazon: trends, projections, and policy implications. Environmental Research Letters 5:014012.

França F, Louzada J, Korasaki V, Griffiths H, Silveira JM, Barlow J. 2016. Do space-for-time assessments underestimate the impacts of logging on tropical biodiversity? An Amazonian case study using dung beetles. Journal of Applied Ecology 53:1098–1105.

Gardner TA et al. 2013. A social and ecological assessment of tropical land uses at multiple scales: the Sustainable Amazon Network. Philosophical Transactions of the Royal Society B: Biological Sciences 368:20120166–20120166.

Gentry AH. 1988. Tree species richness of upper Amazonian forests. Proceedings of the National Academy of Sciences 85:156–159.

Giam X. 2017. Global biodiversity loss from tropical deforestation. Proceedings of the National Academy of Sciences of the United States of America 114:5775–5777. National Acad Sciences.

Gibson L et al. 2011. Primar)-forests are irreplaceable for sustaining tropical biodiversity. Nature 478:378–381. Nature Publishing Group.

Higgins MA, Ruokolainen K, Tuomisto H, Llerena N, Cardenas G, Phillips OL, Vásquez R, Räsänen M. 2011. Geological control of floristic composition in Amazonian forests. Journal of Biogeography 38:2136–2149.

Jenkins CN, Pimm SL, Joppa LN. 2013. Global patterns of terrestrial vertebrate diversity and conservation. Proceedings of the National Academy of Sciences 110:E2602–E2610. National Acad Sciences.

Karp DS, Rominger AJ, Zook j, Ranganathan j, Ehrlich PR, Daily GC. 2012. Intensive agriculture erodes β-diversity at large scales. Ecology Letters 15:963–970.

Karp DS, Ziv G, Zook j, Ehrlich PR, Daily GC. 2011. Resilience and stability in bird guilds across tropical countryside. Proceedings of the National Academy of Sciences 108:21134–21139. National Acad Sciences.

Koh LP, Dunn RR, Sodhi NS, Colwell RK, Proctor HC, Smith VS. 2004. Species coextinctions and the biodiversity crisis. Science 305:1632–1634.

Korasaki V, Braga RE, Zanetti R, Moreira EMS, Vaz-de-Mello EZ, Louzada J. 2013. Conservation value of alternative land-use systems for dung beetles in Amazon: valuing traditional farming practices. Biodiversity and Conservation 22:1485–1499.

Kurz DJ, Nowakowski AJ, Tingley MW, Donnelly MA, Wilcove DS. 2014. Biological Conservation. Biological Conservation 170:246–255. Elsevier Ltd.

Laurance WF et al. 2014. A global strategy for road building. Nature: 1–7. Nature Publishing Group.

Lees AC, Moura NG, de Almeida AS, Vieira ICG. 2015. Poor Prospects for Avian Biodiversity in Amazonian Oil Palm. PloS one 10:e0122432.

MacArthur R, Levins R. 1967. The limiting similarity, convergence, and divergence of coexisting species. American Naturalist 101:377–385. JSTOR.

Mahood SP, Lees AC, Peres CA. 2011. Amazonian countryside habitats provide limited avian conservation value. Biodiversity and Conservation 21:385–405.

Mayfield MM, Daily GC. 2005. Countryside biogeography of neotropical herbaceous and shrubby plants. Ecological Applications 15:423–439. Eco Soc America.

Mäki S, Kalliola R, Vuorinen ?. 2001. Road construction in the Peruvian Amazon: process, causes and consequences. Environmental Conservation 28:199–214. Cambridge Univ Press.

Mendenhall CD, Sekercioglu CH, Brenes E’O, Ehrlich PR, Daily GC. 2011. Predictive model for sustaining biodiversity in tropical countryside. Proceedings of the National Academy of Sciences 108:16313–16316. National Acad Sciences.

Moura N, Lees AC, Andretti CB, Davis BJW, Solar RRC, Aleixo A, Barlow J, Eerriera A, Gardner TA. 2013. Avian biodiversity in multiple-use landscapes of the Brazilian Amazon. Biological Conservation 167:339–348. Elsevier Ltd.

Moura NG, Lees AC, Aleixo A, Barlow J, Berenguer E, Eerreira A, Mac Nally R, Thomson JR, Gardner TA. 2016. Idiosyncratic responses of Amazonian birds to primary forest disturbance. Oecologia 180:903–916. Springer Berlin Heidelberg.

Newbold T, Scharlemann JPW, Butchart SHM, Sekercioglu CH, Joppa L, Alkemade R, Purves DW. 2014. E’unctional traits, land-use change and the structure of present and future bird communities in tropical forests. Global Ecology and Biogeography 23:1073–1084.

Peh KSH, Sodhi NS, de Jong J, Sekercioglu C, Yap CAM, Lim SLH. 2006. Conservation value of degraded habitats for forest birds in southern Peninsular Malaysia. Diversity and Distributions 12:572–581.

Phalan B, Onial M, Balmford A, Green RE. 2011. Reconciling E’ood Production and Biodiversity Conservation: Land Sharing and Land Sparing Compared. Science 333:1289–1291.

Plummer M. 2003. JAGS: A program for analysis of Bayesian graphical models using Gibbs sampling, in K. Homik, E. Leisch, and A. Zeileis, editors. Vienna, Austria.

Pomara LY, Ruokolainen K, Tuomisto H, Young KR. 2012. Avian composition co-varies with floristic composition and soil nutrient concentration in Amazonian upland forests. Biotropica 44:545–553.

Przyborowski J, Wilenski H. 1940. Homogeneity of results in testing samples from Poisson series: With an application to testing clover seed for dodder. Biometrika 31:313–323.

Pulgar Vidal M, Gamboa Moquillaza P. 2013. Reserva Nacional Allpahuayo Mishana Plan Maestro 2013 - 2018. Pages 1–24. Lima, Peru.

Ranganathan J, Daniels RJR, Chandran MDS, Ehrlich PR, Daily GC. 2008. Sustaining biodiversity in ancient tropical countryside. Proceedings of the National Academy of Sciences 105:17852–17854. National Acad Sciences.

Ravikumar A, Sears RR, Cronkleton P, Menton M, Pérez-Ojeda del Arco M. 2017. Is small-scale agriculture really the main driver of deforestation in the Peruvian Amazon? Moving beyond the prevailing narrative. Conservation Letters 10:170–177.

Remsen JV Jr, Parker TA 111. 1983. Contribution of river-created habitats to bird species richness in Amazonia. Biotropica:223–231. JSTOR.

Rosenberg GH. 1990. Habitat specialization and foraging behavior by birds of Amazonian river islands in northeastern Peru. The Condor:427–443. JSTOR.

Sekercioglu C, Loarie SR, Oviedo Brenes l·’, Ehrlich PR, Daily GC. 2007. Persistence of forest birds in the Costa Rican agricultural countryside. Conservation Biology 21:482–494.

Socolar J, Robinson SK, Terborgh JW. 2013. Bird Diversity and Occurrence of Bamboo Specialists in Two Bamboo Die-Offs in Southeastern Peru. The Condor 115:253–262.

Socolar JB, Gilroy JJ, Kunin WE, Edwards DP. 2016. How should beta-diversity inform biodiversity conservation? Trends in Ecology & Evolution 31:67–80. Elsevier Ltd.

Solar RR de C et al. 2015. How pervasive is biotic homogenization in human-modified tropical forest landscapes? Ecology Letters 18:1108–1118.

Steege ter H et al. 2013. Hyperdominance in the Amazonian tree flora. Science 342:1243092–1243092.

Tuomisto H, Ruokolainen K, Kalliola R, Linna A, Danjoy W, Rodriguez Z. 1995. Dissecting Amazonian biodiversity. Science 269:63–66.

Tyukavina A, Hansen MC, Potapov PV, Krylov AM, Goetz SJ. 2015. Pan-tropical hinterland forests: mapping minimally disturbed forests. Global Ecology and Biogeography 25:151–163.

Wittmann F, Junk WJ, Piedade MTF. 2004. The várzea forests in Amazonia: flooding and the highly dynamic geomorphology interact with natural forest succession. Forest Ecology and Management 196:199–212.

Wittmann F, Schongart J, Montero JC, Motzer T, Junk WJ, Piedade MTF, Queiroz HL, Worbes M. 2006. Tree species composition and diversity gradients in white-water forests across the Amazon Basin. Journal of Biogeography 33:1334–1347.

